# *Muribaculaceae* genomes assembled from metagenomes suggest genetic drivers of differential response to acarbose treatment in mice

**DOI:** 10.1101/2020.07.01.183202

**Authors:** Byron J. Smith, Richard A. Miller, Thomas M. Schmidt

## Abstract

The drug acarbose is used to treat diabetes, and, by inhibiting α-amylase in the small intestine, increases the amount of starch entering the lower digestive tract. This results in changes to the composition of the microbiota and their fermentation products. Acarbose also increases longevity in mice, an effect that has been correlated with increased production of the short-chain fatty acids propionate and butyrate. In experiments replicated across three study sites, two distantly related species in the bacterial family *Muribaculaceae* were dramatically more abundant in acarbose-treated mice, distinguishing these responders from other members of the family. Bacteria in the family *Muribaculaceae* are predicted to produce propionate as a fermentation end product and are abundant and diverse in the guts of mice, although few isolates are available. We reconstructed genomes from metagenomes (MAGs) for nine populations of *Muribaculaceae* to examine factors that distinguish species that respond positively to acarbose. We found two closely related MAGs (B1A and B1B) from one responsive species that both contain a polysaccharide utilization locus with a predicted extracellular α-amylase. These genomes also shared a periplasmic neopullulanase with another, distantly related MAG (B2) representative of the only other responsive species. This gene differentiated these three MAGs from MAGs representative of non-responding species. Differential gene content in B1A and B1B may be associated with the inconsistent response of this species to acarbose across study sites. This work demonstrates the utility of culture-free genomics for inferring the ecological roles of gut bacteria including their response to pharmaceutical perturbations.

**Importance:** The drug acarbose is used to treat diabetes by preventing the breakdown of starch in the small intestine, resulting in dramatic changes in the abundance of some members of the gut microbiome and its fermentation products. In mice, several of the bacteria that respond most positively are classified in the family *Muribaculaceae*, members of which produce propionate as a primary fermentation product. Propionate has been associated with gut health and increased longevity in mice. We found that genomes of the most responsive *Muribaculaceae* showed signs of specialization for starch fermentation, presumably providing them a competitive advantage in the large intestine of animals consuming acarbose. Comparisons among genomes enhance existing models for the ecological niches occupied by members of this family. In addition, genes encoding one type of enzyme known to participate in starch breakdown were found in all three genomes from responding species, but none of the other genomes.

## Introduction

The mammalian gut microbiome is a complex ecological system that influences energy balance (1), pathogen resistance (2), and inflammation (3), among other processes with importance to host health. Understanding how the bacterial inhabitants of the gut respond to pharmaceutical and dietary perturbations is a major step in developing a predictive framework for microbiome-based therapies. Acarbose (ACA) is an α-glucosidase inhibitor prescribed for the treatment of type 2 diabetes mellitus because it reduces the absorption of glucose from starch in the small intestine (4). In rats, ACA has been shown to increase the amount of starch entering the lower digestive system after a meal (5). ACA treatment also changes the composition of the gut microbiota and its fermentation products in many rodents (5–14). Interestingly, long-term treatment with ACA has been shown to substantially increase longevity in male mice and to a lesser extent in females (15–17).

Previously we found that the relative abundance of multiple bacterial species as well as the concentrations of propionate and butyrate respond to long term treatment with ACA (14). That study was notable in being replicated across three sites: The University of Michigan (UM) in Ann Arbor, The University of Texas Health Science Center at San Antonio (UT), and The Jackson Laboratory (TJL) in Bar Harbor, Maine. At UM and TJL one highly abundant bacterial species was enriched nearly 4-fold in ACA-treated mice. This species, defined at a 97% identity threshold of the 16S rRNA gene V4 region and designated as OTU-1, was classified as a member of the family *Muribaculaceae* in the order Bacteroidales. OTU-1 was also present and abundant at UT but was not significantly more abundant in ACA-treated mice relative to controls. Instead, a different *Muribaculaceae* species, designated OTU-4, was found to be highly abundant and enriched 4-fold in ACA-treated mice, but was nearly absent at UM and TJL. Other *Muribaculaceae* were also identified as among the most abundant members of the mouse gut microbiota across the three sites, although none of these were found to be enriched during ACA treatment.

Members of the family *Muribaculaceae*—previously referred to as family S24-7 after an early clone (18, 19), or sometimes as *Candidatus* Homeothermaceae (20)—have only recently been isolated (21–23) despite being a common and abundant inhabitant of the mammalian gut, especially in mice (20, 22). Studies using culture-free methods suggest that the *Muribaculaceae* specialize on the fermentation of complex polysaccharides (20, 22), much like members of the genus *Bacteroides*, which is also a member of the order Bacteroidales. Genomic analysis has also suggested that the capacity for propionate production is widespread in the family (20). Recently, techniques have been developed to reconstruct genomes of uncultivated members of bacterial communities (24, 25). Based on 30 such metagenome assembled genomes (MAGs) they reconstructed, Ormerod and colleagues (20) proposed that the *Muribaculaceae* fall into three distinct carbohydrate utilization guilds, which they describe as specialists on α-glucans, plant glycans, and host glycans, respectively. While it is reasonable to expect that α-glucan specialists would benefit the most from the large influx of starch to the gut resulting from ACA treatment, this prediction has not been tested, and physiological inferences based on the genome content of members of the family have been largely divorced from biological observations.

Experimental perturbations of complex microbial communities present an opportunity to observe ecological features of many bacterial taxa without cultivated members and generate hypotheses about their physiology. Given the observed, dramatically increased relative abundance of OTU-1 and OTU-4 (here referred to as “responders”) in mice treated with ACA, we hypothesize that these species are capable of robust growth on starch, while the other *Muribaculaceae* found in the study (“non-responders”), lack the genomic features necessary for the utilization of polysaccharides that reach the colon in greater quantities following ACA treatment.

Alternatively, responders may be resistant to the inhibitory effects of ACA, or benefit from elevated levels of intermediate starch degradation products. Since isolates of the *Muribaculaceae* strains in these mice are not available for characterization, we took a comparative genomic approach to explore their functional potential.

Most of the research on the genomic components of polysaccharide degradation in gram negative bacteria has been carried out in the genus *Bacteroides*, and in particular *B. thetaiotaomicron* (26). Starch utilization in *B. thetaiotaomicron* is dependent on an ensemble of eight proteins, SusRABCDEFG that enable recognition, binding, hydrolysis, and import of starch and related polysaccharides (27). Homologs of SusC and SusD characterize all known polysaccharide utilization systems in this clade (28), are encoded in Sus-like genomic regions known as polysaccharide utilization loci (PULs), and are widespread in the phylum Bacteroidetes (29). The molecular range of these systems is determined by the carbohydrate-active enzymes and structural proteins they encode, based on the specificity of glycoside hydrolase (GH) and carbohydrate binding module (CBM) domains, which have been extensively cataloged in the CAZy and dbCAN databases (30–33).

Here MAGs from the feces of mice at UT and UM are analyzed in the context of previously generated MAGs and cultivar genomes to explore two closely related questions about the niche of OTU-1 and OTU-4 in the lower digestive system. First, why do these species each increase in relative abundance with ACA treatment, while other species of *Muribaculaceae* do not? And second, why is the response of OTU-1 site specific? Despite similar patterns of abundance at their respective sites, the two responding species seem to be only distantly related, sharing just 90% of nucleotides in their 16S rRNA gene V4 hypervariable region (14). We nonetheless find genomic evidence that OTU-1 and OTU-4 occupy overlapping niches, specializing in the degradation of α-glucans, a role not held by the other *Muribaculaceae* described in this study. In addition, we identify two distinct genomic variants of OTU-1, referred to as B1A and B1B, which are differentially distributed between UM and UT and have functionally relevant differences in gene content.

Reconstructing genomes from metagenomes allows for the comparison of the functional potential of *Muribaculaceae* at UM and UT. This work demonstrates the utility of culture-free genomics to understand the potential ecological roles of these key members of the mouse gut microbial community and explore several hypotheses that may explain differences in the distribution and response of these bacteria to ACA treatment. Hypotheses derived from this analysis provide a foundation for future physiological studies in recently obtained cultivars. While a large fraction of host-associated bacterial species are without isolated representatives (34), let alone characterized (35), combining experimental data from complex communities with the analysis of reconstructed genomes provides a powerful tool for expanding understanding to these understudied taxa.

## Results

### Recovered population genomes are of high quality and resemble other *Muribaculaceae* genomes

MAGs were constructed for 9 populations classified as members of the family *Muribaculaceae*, including for two species, OTU-1 and OTU-4, previously shown to respond positively to ACA. All 9 novel MAGs are estimated to be more than 84% complete and all had less than 2% estimated contamination based on the recovery of ubiquitous, single-copy genes (Tbl. 1). The median N50 statistic was approximately 61 kbp, suggesting that assembly was suitable for inferring the genomic context of functional genes. For OTU-1, two closely related genomic variants were recovered, here designated B1A and B1B, possessing 0.56 and 0.31 Mbp of unshared sequence, respectively (Tbl. 2). We designate the MAG constructed for OTU-4 as B2. MAGs obtained from non-responding species are designated B3 through B8. Estimated genome sizes, %GC, and number of predicted genes are all similar to reference genomes from cultivated members of the family *Muribaculaceae*.

To confirm the assertion that each of the reconstructed genomes is representative of a previously described *Muribaculaceae* species identified in these mice (14), the median mapping rate of metagenomic reads to protein coding features for each MAG was compared to the relative abundance of the associated 16S rRNA gene across matched amplicon libraries. Reassuringly, cosine similarities were above 0.88 for all MAGs, suggesting robust concordance in coverage between the shotgun metagenomic and amplicon libraries. Correlated coverage statistics can be found in the Supplementary Results (see `build/otu_correlation_and_aca_response.ipynb.html` at https://doi.org/10.5281/zenodo.4450697).

**Table 1:** *Summary of novel MAGs, previously described, high-quality MAGs, and cultivar genomes*: Muribaculum intestinale *(Mi)*, Duncaniella muris *(Dm)*, Duncaniella freteri *(Df)*, Duncaniella dubosii *(Dd)*, Paramuribaculum intestinale *(Pi)*, Candidatus *Homeothermus arabinoxylanisolvens (Ha), Candidatus Amulumruptor caecigallinarius (Ac). *’s indicate MAGs from responder populations*.

| Genome | CheckM Completeness | Scaffolds | Length (Mbp) | N50 | GC | in (14) |
| --- | --- | --- | --- | --- | --- | --- |
| B1A* | 95% | 286 | 3.1 | 23,337 | 46.6% | OTU-1 |
| B1B* | 94% | 320 | 2.8 | 19,144 | 47.0% | OTU-1 |
| B2* | 96% | 116 | 2.7 | 75,014 | 50.6% | OTU-4 |
| B3 | 93% | 62 | 2.7 | 80,587 | 55.8% | OTU-8 |
| B4 | 97% | 30 | 2.7 | 127,141 | 55.4% | OTU-5 |
| B5 | 89% | 124 | 2.3 | 46,359 | 54.2% | OTU-6 |
| B6 | 90% | 87 | 2.3 | 61,151 | 54.2% | OTU-39 |
| B7 | 84% | 217 | 2.9 | 78,457 | 48.0% | OTU-30 |
| B8 | 87% | 154 | 2.2 | 33,525 | 53.4% | OTU-11 |
| Mi | 99% | 1 | 3.3 | 3,307,069 | 50.1% | - |
| Dm | 99% | 80 | 3.3 | 93,062 | 50.8% | - |
| Df | 99% | 5 | 3.6 | 2,271,962 | 48.5% | - |
| Dd | 97% | 5 | 3.7 | 3,598,157 | 47.9% | - |
| Pi | 98% | 378 | 2.9 | 13,720 | 53.1% | - |
| Ha | 99% | 60 | 2.8 | 80,867 | 50.3% | - |
| Ac | 90% | 157 | 2.4 | 41,191 | 51.9% | - |

**Table 2:** Summary of variant specific features in two highly similar MAGs

|  | B1A |  | B1B |  |
| --- | --- | --- | --- | --- |
|  | Total | Specific | Total | Specific |
| Nucleotide length (Mbp) | 3.07 | 0.56 | 2.82 | 0.31 |
| Genes | 2,612 | 307 | 2,419 | 291 |
| OPFs (unique) | 2,221 | 242 | 2,096 | 234 |
| KOs (unique) | 1,026 | 64 | 1,000 | 44 |
| COGs (unique) | 539 | 15 | 528 | 8 |
| Pfam domains (unique) | 3,657 | 635 | 3,490 | 488 |
| CAZy domains (unique) | 252 | 35 | 240 | 18 |

### Phylogenetics

To better understand the evolutionary relationships between these organisms, a concatenated gene tree (Fig. 1 panel A and Supplementary Fig. S1 at https://doi.org/10.5281/zenodo.4450697) was constructed for the 9 new MAGs along with publicly available MAGs and isolate genomes (20–23) The tree was rooted by five other Bacteroidales species: *Bacteroides ovatus* (ATCC-8483), *Bacteroides thetaiotaomicron* VPI-5482, *Porphyromonas gingivalis* (ATCC-33277), *Barnesiella viscericola* (DSM-18177), and *Barnesiella intestinihominis* (YIT-11860). Most internal nodes were supported with high topological confidence (>95% bootstrap support), and the placement of the MAGs derived by Ormerod and colleagues (20) was highly consistent with their published tree. To further check the robustness of our phylogeny, a second, approximate maximum likelihood tree was constructed based on the *rpoB* gene, which is generally not thought to be transmitted horizontally (with exceptions (36)), While *rpoB* was not annotated in a number of the genomes, and some nodes were unable to be resolved, this approach largely confirmed the topology of the concatenated gene tree (see Supplementary Fig. S2 at https://doi.org/10.5281/zenodo.4450697). The estimated phylogeny shows that the newly generated MAGs encompass most of the documented diversity of *Muribaculaceae* (Fig. 1A). While many of the novel MAGs are phylogenetically similar to previously described genomes, two of the MAGs, B3 and B4, are notably diverged from the most closely related taxa. This demonstrates, that despite a growing number of *Muribaculaceae* genomes deposited in public repositories, novel taxa remain to be described.

**Figure 1:**
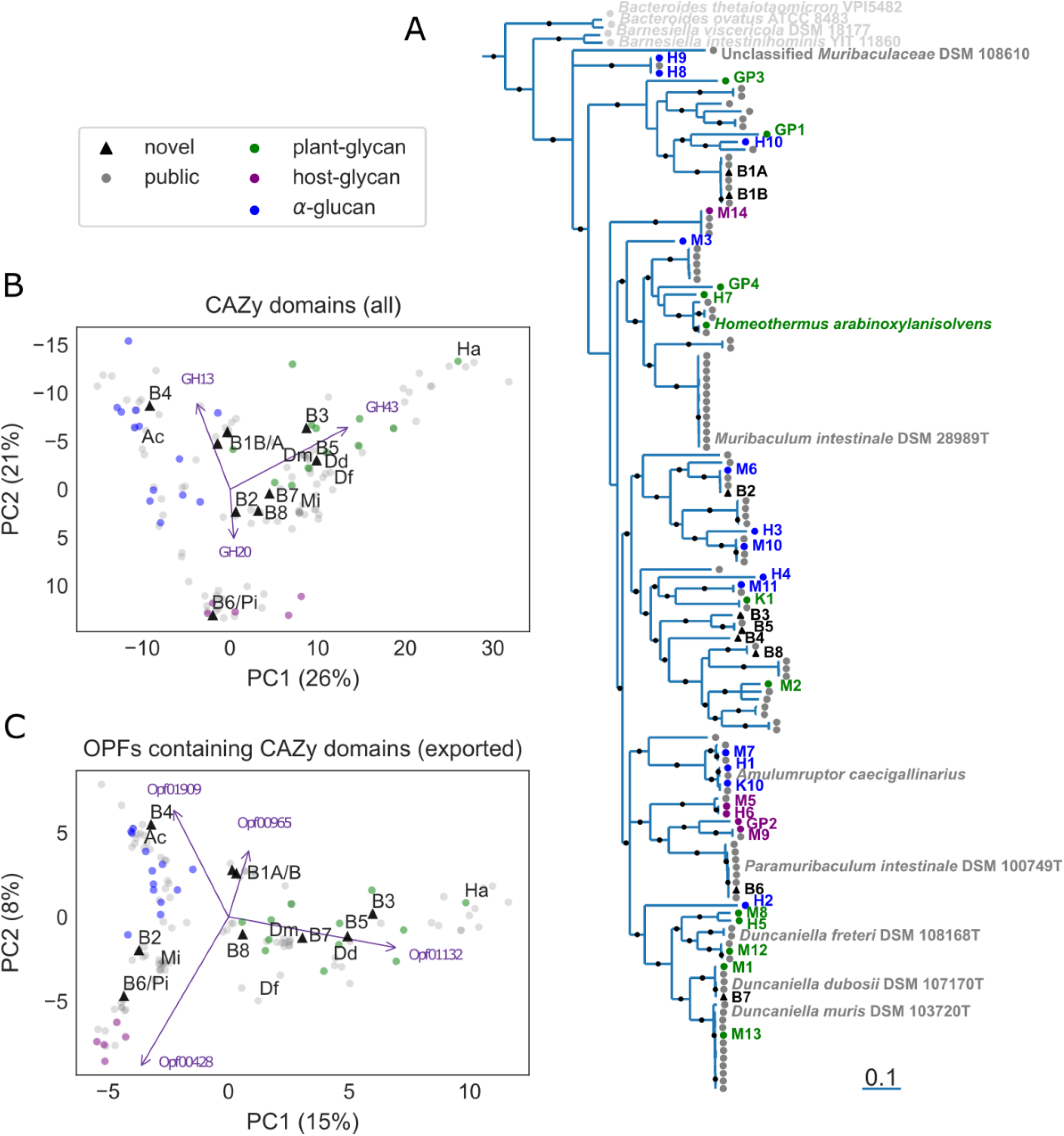
Comparison of novel and previously described Muribaculaceae genomes. Novel MAGs (labeled “B1A” through “B8”) are combined with publicly available genomes and MAGs, as well as 30 MAGs constructed in (20) that are hypothesized to reflect three polysaccharide utilization guilds (marker colors). (**A**) MAGs reconstructed in this study (highlighted in black) are placed in a phylogenetic context using an approximate maximum-likelihood concatenated gene tree based on an amino acid alignment of 36 shared, single-copy genes. The tree is rooted by Porphyromonas gingivalis ATCC-33277 (not shown) and four additional Bacteroidales genomes are included as an outgroup. Nodes with less than 70% confidence are collapsed into polytomies and topological support greater than 95% is indicated (black dots on internal branches). Branch lengths indicate an estimate of expected substitutions per site. A version of this panel with GenBank accessions for all publicly available genomes is available as Supplementary Fig. S1 at https://doi.org/10.5281/zenodo.4450697. Functional comparisons are visualized (**B, C**) by plotting the first two principal components (PCs) of an ordination based on counts of predicted proteins annotated with GH and CBM domains either (**B**) aggregated by CAZy family or (**C**) possessing a signal peptide and aggregated by OPF. (**B, C**) Purple arrows indicate the contributions of the labeled features, and axes are labeled with the fraction of all variation accounted for by that PC. Novel MAGs (black triangles) are labeled, as are 7 previously described cultivar genomes and high-quality MAGs: Candidatus Homeothermus arabinoxylanisolvens (Ha), Muribaculum intestinale (Mi), Duncaniella muris (Dm), Duncaniella freteri (Df), Duncaniella dubosii (Dd) Paramuribaculum intestinale (Pi), Candidatus Amulumruptor caecigallinarius (Ac).

### Novel protein families

Annotations based on alignment to a database of previously characterized sequences may provide only limited insight into the function of gene products, in particular for genes from largely unstudied families of bacteria. In order to identify previously uncharacterized orthologous groups of genes, *de novo* clustering (37) was carried out based on amino acid similarity of all 348,908 putative protein coding genes found in the 9 novel MAGs, 30 MAGs previously generated by Ormerod and colleagues (20), all 98 *Muribaculaceae* genome assemblies available from GenBank as of this work, and 5 reference genomes from other members of the order Bacteroidales. The resulting 16,859 clusters with more than one member contain 315,581 sequences (90%) and are referred to here as operational protein families (OPFs). While a fraction of these clusters may be due to spurious sequence similarity and without biological relevance, 12,876 have representatives in at least three genomes, increasing the likelihood that these reflect evolutionarily conserved protein sequences. Of this higher confidence set, only 4,448 have members annotated with any COG, KO, or putative function. The remaining 8,528 unannotated OPFs encompass 111,177 predicted protein sequences across the 142 genomes.

To better understand the relationship between our *de novo* clusters and previously described annotations, we inspected the concordance between known GH13 subfamilies, which possess α-amylase activities on different starches (38), and the 20 OPFs with these domains (see Supplementary Results `build/gh13_families_to_opfs_mapping.ipynb.html` at https://doi.org/10.5281/zenodo.4450697). We find that most OPFs are composed of members with just one subfamily, suggesting that our clusters are consistent with these known groupings. Several OPFs are dominated by members without a more specific subfamily, and while outside the scope of this study, these may offer hypotheses for additional divisions within GH13. We provide an estimated phylogenetic tree of all GH13 domains, their OPF assignments, and their predicted localization in the Supplementary Data `build/gh13_tree.nwk` at https://doi.org/10.5281/zenodo.4450697. Detailed annotations of predicted genes in MAG and reference genomes with OPFs, COGs, KOs, and Pfam and CAZy domains are available as Supplementary Tbl. S1 at https://doi.org/10.5281/zenodo.4450697.

### Ordination of gene content

To better understand the polysaccharide utilization potential encoded by the novel MAGs and other available *Muribaculaceae* genomes, we performed an ordination of the counts of genes with predicted homology to GH or CBM domains (see Fig. 1 panel B). This holistic analysis of genes that could plausibly be involved in polysaccharide degradation recapitulates the three clusters previously described by Ormerod and colleagues for the first 30 MAGs from the family (20), including associations of the hypothesized α-glucan, plant glycan, and host glycan guilds with GH13, GH43, and GH20 domains, respectively. However, given the ambiguous, intermediate placement of many newer genomes, it also suggests exceptions to the three-guild model. Notably, none of B1A, B1B, or B2—MAGs representative of responding species—were placed cleanly within the α-glucan guild as we had hypothesized.

It is likely that a more nuanced approach to comparing *Muribaculaceae* genomes will yield enhanced predictions of functional capacities. To better reflect *a priori* assumptions about the cellular mechanisms of polysaccharide utilization in the family, the same analysis was repeated, but with two modifications. First, since proteins must be localized to the cell envelope in order to participate in the breakdown of large, extracellular polysaccharides, genes were only counted if they include an N-terminal signal peptide sequence enabling export from the cytoplasm. Second, given the potential for OPFs to reflect orthologous functions better than domain level homology, the OPF designations were tallied for predicted proteins with homology to GHs and CBMs, rather than the domains themselves. While this more targeted analysis illustrates similar trends to before, the distinction between the three clusters appears visually more defined (Fig. 1 panel C). Interestingly, in this ordination B1A, B1B and several closely related genomes now occupy a space proximate to but distinct from the previously hypothesized α-glucan guild.

Reassuringly, OPFs driving the separation between clusters possess domains matching the original guild descriptions (20). For instance, Opf00428 is associated with MAGs previously assigned to the host glycan guild and all but one member of this protein family include regions homologous to GH20. Likewise, Opf01132 parallels the plant glycan guild and all predicted proteins in this family include a GH43 domain. Surprisingly, not all protein families sharing the same domains are equivalent. Although Opf01909 and Opf00965 both possess a predicted GH13 domain—and are the first and fourth most positively weighted features in PC2, indicating an association with the α-glucan guild—the latter is also enriched in the plant glycan guild while the former is not; of the 12 MAGs originally classified to the plant glycan guild, 11 have at least one copy of Opf00965 with a signal peptide versus just one MAG with Opf01909. Based on annotation by the dbCAN2 meta server (32), The GH13 domain in Opf01909 members is classified to the recently defined subfamily 42 (39), however Opf00965 is not assigned to a more precise subfamily (see Supplementary Results `build/gh13_families_to_opfs_mapping.ipynb.html` at https://doi.org/10.5281/zenodo.4450697). While a detailed enrichment analysis is outside the scope of this work, this suggests that OPFs may indeed provide more functional resolution than annotations based on existing reference databases.

### Analysis of MAGs from species responsive to ACA treatment suggests genes involved in starch utilization

Based on the characterization of genes and genomic regions with a role in starch utilization in the closely related genus *Bacteroides*, it is plausible that an α-amylase localized to the outer membrane (OM) may be common to starch utilizing bacteria in the order Bacteroidales (40). Indeed, B1A and B1B both have three genes predicted to code for GH13 containing, OM-localized lipoproteins (B1A_01660, B1A_01692, B1A_02267 in B1A and B1B_01504, B1B_01538, B1B_02118 in B1B), each in a separate PUL (see Fig. 2). While it also includes members without this activity, GH13 is the most common family of α-amylases (41). These genomic regions also possess additional genes with carbohydrate-active domains that are expected to interact with α-glucans.

**Figure 2:**
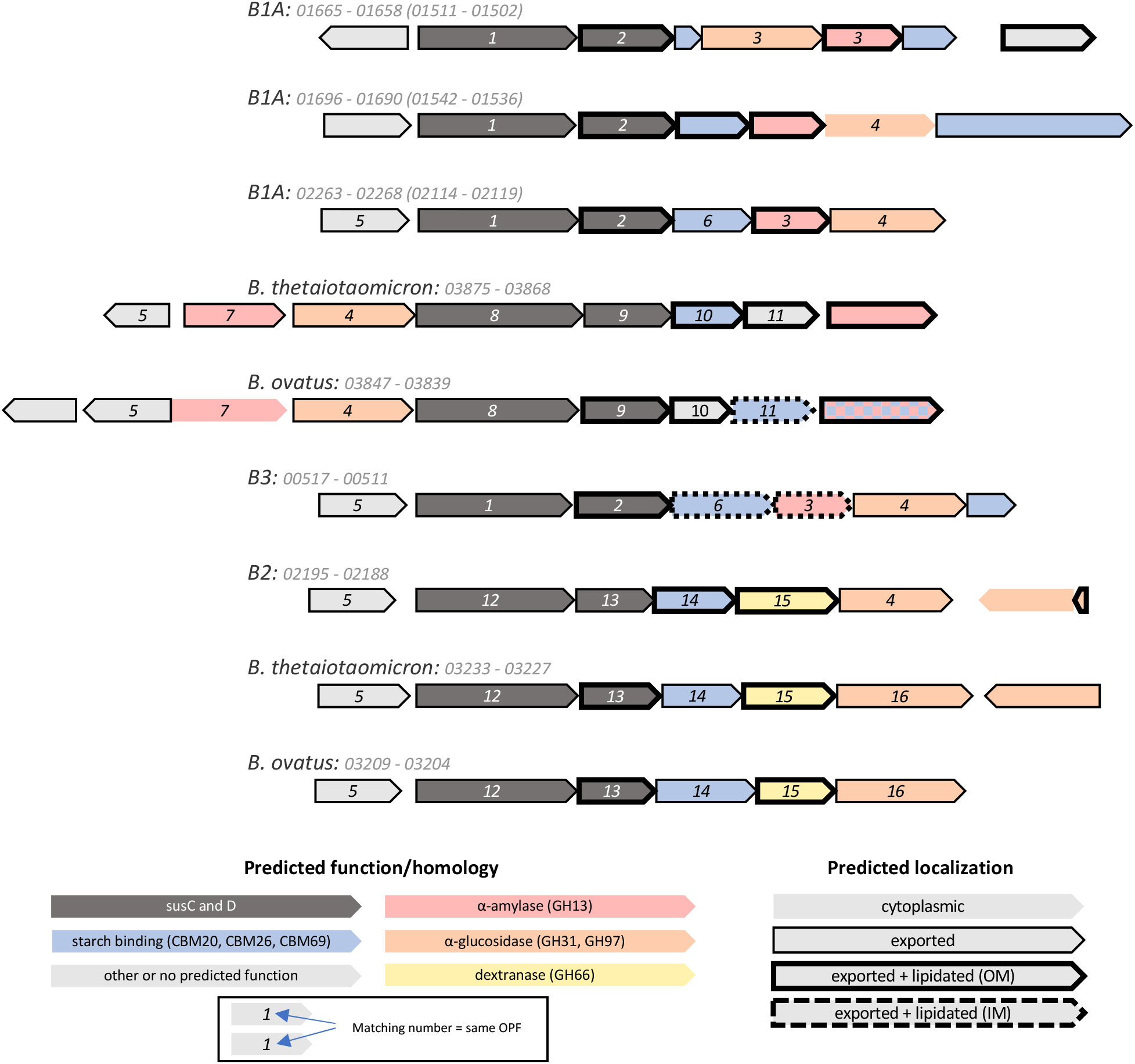
Diagrams of PULs plausibly reflecting activity on starch and/or dextran in B1A, B2, B3, B. thetaiotaomicron, and B. ovatus. Regions are labeled with the genome name and the interval of gene numbers. For B1A PULs, the matching gene numbers in B1B are noted in parentheses. ORFs are depicted as arrows pointed 5’ to 3’ along the coding sequence, and colors indicate homology to genes and domains known to participate in either starch or dextran utilization. ORF outlines indicate predicted localization based on the presence of an N-terminal signal peptide and nearby residues. Matching numbers indicate homology based on OPF clustering, and are arbitrarily assigned. (1: Opf01209, 2: Opf02007, 3: Opf02000, 4: Opf00042, 5: Opf01405, 6: Opf02584, 7: Opf01765, 8: Opf00431, 9: Opf09589, 10: Opf15294, 11: Opf14773, 12: Opf01209, 13: Opf03138, 14: Opf04347, 15: Opf04327, 16: Opf16791).

Besides B1A and B1B, B3 is the only other MAG to possess a putative PUL coding for a full complement of predicted starch-active proteins. This PUL has a large degree of synteny with one of the three matching PULs found in both B1A and B1B, and also includes two OPFs with members also found in the Sus-operon of *B. thetaiotaomicron*, suggesting shared function. However, while B3 also has a GH13 containing lipoprotein (B3_00513), its predicted localization is to the inner membrane, not the OM. It is unclear whether this explains B3’s non-response in ACA-treated mice. Only two other IM-localized GH13 containing proteins are found across all of the genomes analyzed here (Supplementary Data `build/gh13_tree.nwk` at https://doi.org/10.5281/zenodo.4450697). Plausible OM-localized, GH13 containing proteins are not found in any non-responders. While this characteristic does not seem to perfectly discriminate responder from non-responders—B2 also lacks such a gene—it nonetheless demonstrates concordance between inferred genomic features and observed population dynamics of the corresponding species.

Despite the absence of a GH13 domain on the OM, it is plausible that B2 is capable of degrading starch using other enzymatic machinery. We speculate about one putative locus (Fig. 2), which is highly syntenic with characterized dextran PULs in *B. thetaiotaomicron* and *B. ovatus* (42–44). While the only predicted, OM-localized GH contains a GH66 domain (B2_02191), other proximal genes have domains with known affinity for starch, including GH97, CBM20, and CBM69.

To expand the search for relevant genetic features, *de novo* protein clusters were filtered to those with members in B1A and B2, and in at most one of the non-responders, B3-B8. Of these 52 OPFs, two stood out as particularly relevant. Opf01405 includes SusR, the regulator of transcription of the starch utilization system in *B. thetaiotaomicron*, as well as its homolog in *B. ovatus*. It is an apparent subcluster of the larger family defined by K21557, and in many cases is encoded directly upstream of *susC* in putative PULs that are considered likely to have affinity for α-glucans. In each of B1A and B1B, one of the three putative starch PULs encodes a member of Opf01405, and it is similarly located in PULs with starch-active CBM and GH domains in B2 and B3. Interestingly, this OPF is not found in any of the other novel MAGs. In addition, of the seven MAGs constructed by Ormerod *et al*. that encode a member of this cluster, five of them are classified to the α-glucan guild. It is plausible that members of Opf01405 share a functional role regulating transcriptional responses to α-glucans.

Finally, we find one example of a GH13 containing OPF that is found in MAGs representing the responder, B1A, B1B, and B2, but none of the other MAGs generated in this study. Opf01765, which recapitulates K21575, includes SusA: the periplasmic neopullulanase of *B. thetaiotaomicron* and an important component of starch utilization in that organism (45). What’s more, the OPF is found in twelve of the thirteen α-glucan and a minority of the plant glycan guild members. Interestingly, although it is encoded by the Sus operon in *B. thetaiotaomicron* and its homologous locus in *B. ovatus*, in the *Muribaculaceae* members of Opf01765 are not encoded in PULs, with only 7 of 44 members within 25 kbp of the closest PUL *susC* homolog, and all of these either upstream of *susC* or on the opposite strand.

### Unshared gene content in B1A and B1B

Two distinct genomic variants were associated with OTU-1 with one found in a majority of the UT mouse metagenomes, and the other ubiquitous at UM. Using the QUAST package (46), 18.1% of the B1A MAG sequence and 11.0% of B1B were found to not align to the other (Tbl. 2). While these hundreds of kbp may in part reflect errors in genome recovery, much of the unaligned length suggests differences in gene content between these populations of OTU-1. This observation was confirmed by assessing the mapping of metagenomic reads against predicted protein coding genes in each variant (Fig. 3). For each pairing of metagenomic read library to genomic variant, gene coverage was normalized by the median gene coverage in order to identify genes with conspicuously fewer reads in particular subsets of the mice. Metagenomic libraries were manually chosen as unambiguous representatives of either B1A or B1B based on concordance in their coverage profiles (see Fig. 3) and these were used to systematically identify genes differentiating the two populations.

**Figure 3:**
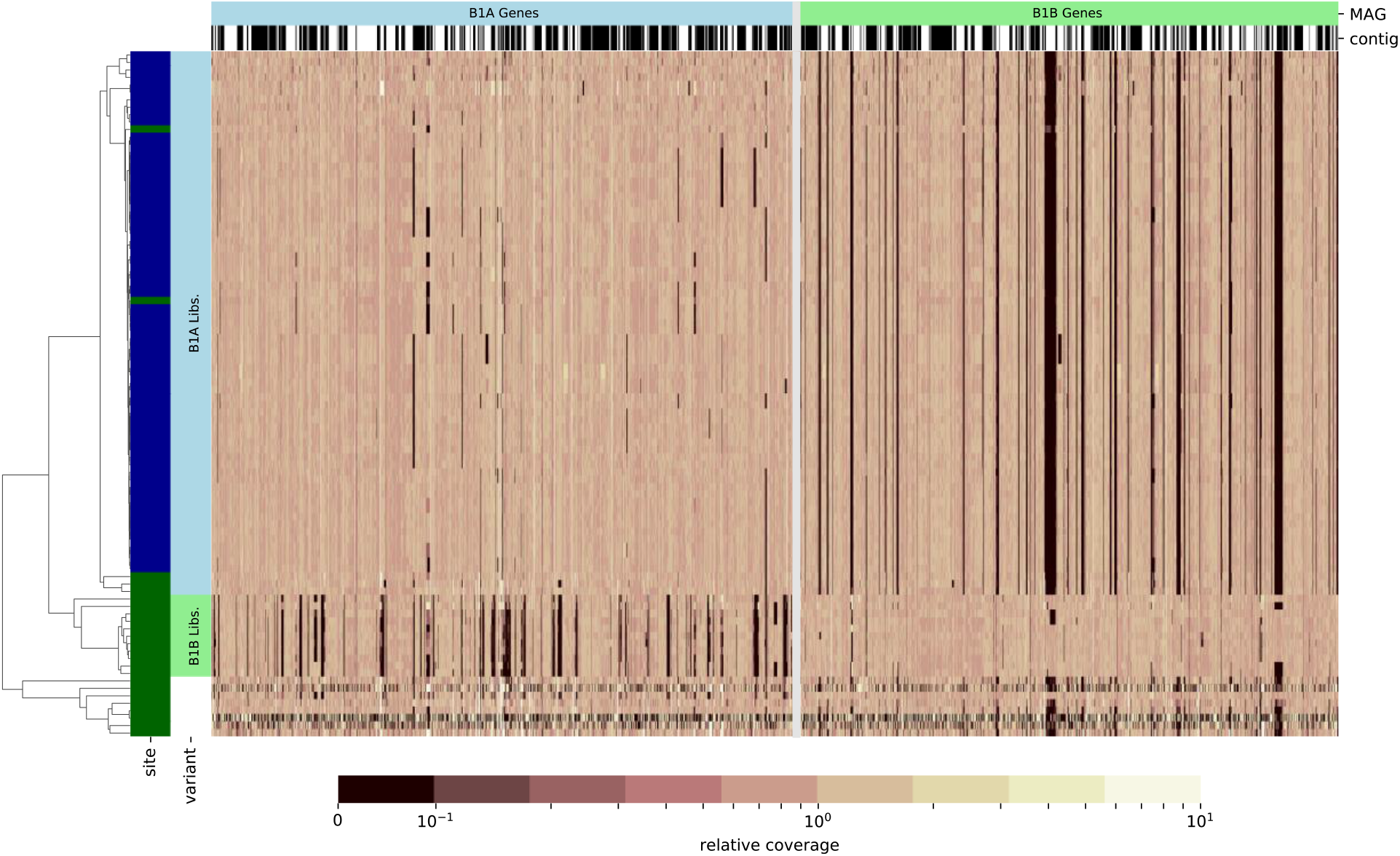
Visualization of differential gene content in two OTU-1 populations. Heatmaps depict relative mapping coverage of metagenomes against putative protein coding genes in MAGs B1A (left of grey line) or B1B (right). Rows represent one or more pooled libraries for each mouse included in the study and columns represent individual genes. Alternating black and white spans over heatmap columns indicate contig boundaries in each MAG; the orientation and order of contigs is arbitrary. All coverage values are normalized to the median coverage of that genome’s features within each mouse. The site at which each mouse was housed is indicated by colored spans on the left (UT: dark green, UM: dark blue), and mice identified as unambiguous representations of each population are indicated (B1A: light blue, B1B: light green, uncertain: white). Rows are ordered based on a hierarchical clustering by cosine distance, depicted in the tree on the left.

The median normalized mapping depths in each set of libraries against predicted genes in each MAG were compared, providing a measure of the relative enrichment or depletion of genomic sequences between the two populations of OTU-1. Libraries specific to each variant have low coverage over large portions of either the B1A or B1B MAG, suggesting that mice are primarily colonized by one of the two variants, and that a portion of genes are variant specific. At a 5-fold depletion cutoff—selected as a compromise between sensitivity and specificity in identifying genomic regions that differentiate the two—we found 11.7% of predicted genes in B1A were depleted in B1B populations, and 12.0% the reverse (Tbl. 2). Coverage ratios over all genes in B1A and B1B can be found in Supplementary Tbl. S2 (available at https://doi.org/10.5281/zenodo.4450697). While the observed coverage ratios could indicate variation in copy number, differential gene content between variants is a more parsimonious explanation for most loci. These predicted genes reflect 6.2% of unique KOs in B1A and 4.4% in B1B. Interestingly, the fraction of variant specific OPFs is greater, 10.8% and 11.1% respectively, suggesting that *de novo* clustering could be more sensitive to potential differences in physiology.

Given the observation that the relative abundance of OTU-1 was dramatically increased with ACA treatment at UM, while not being significantly affected at UT, and that B1B was not found in metagenomes at UM, we hypothesized that differences in the genomic potential of B1A and B1B could explain the different response to ACA at the two sites.

Genomic regions apparently specific to B1A—defined as an at least 5-fold enrichment in B1A specific libraries relative to B1B specific libraries—include just one PUL (SusC-homolog encoded by B1A_02041). This locus includes a predicted OM-localized, GH30-containing protein. Proteins that contain a GH30 domain have β-glucosylceramidase, β-1,6-glucanase, or β-xylosidase activity (47). Given that this PUL also encodes a periplasmic, GH3 containing protein, it appears to be unlikely that it has specificity for starch. Both variants also possess numerous phage insertions not seen in the other. Likewise, a CRISPR operon (Cas9 encoded by B1B_00401) appears to be specific to B1B.

Most strikingly, a 15 kbp region specific to B1A (from B1A_01550 to B1A_01566) was found to contain numerous genes with homology to cell capsule and exopolysaccharide synthesizing enzymes. Based on annotations with KEGG orthologous groups, these include homologs of *tuaG* (K16698), *tagE* (K00712), *gmhB* (K03273), *gmhA*/*lpcA* (K03271), *hddA* (K07031), *exoO* (K16555), *waaH* (K19354), and *tagF* (K09809). Interestingly, the B1B MAG contains a different such region of about 6.2 kbp (B1B_00746 to B1B_00751) with *wfeD* (K21364), *pglJ* (K17248), and *epsH* (K19425). For each, several of the OPFs in the respective regions were not found anywhere in the opposing genome, suggesting that the makeup of each variant’s exterior surface might be distinctly different.

## Discussion

Mice are a key model system for the study of the mammalian gut microbiome, with an outsized importance in testing mechanistic hypotheses about the roles of this community in host health (48). The generalizability of observations made in mice is a constant concern (48), in part due to extensive difference in taxonomic composition compared to humans (21). Bacteria classified in the family *Muribaculaceae* are abundant in the murine gut microbiome (20). While these bacteria are also found in humans (although at lower abundance), only a few members of this clade have been cultivated and described (21–23). As a result, the ecological roles of these bacteria have not yet been characterized, limiting the value of the mouse as a model system. Better understanding the ecology of *Muribaculaceae* in the murine gut will increase the transferability of microbiome findings from mice to humans. Attempts to study these organisms make use of genomes reconstructed from metagenomic reads, and have suggested—in the absence of experimental data—that members of the family consume a diversity of polysaccharides in the lower gut.

Here we have extended that approach to nine new genomes, and associated those with species for which changes in relative abundance in response to ACA treatment have been experimentally assessed. This enabled us to explore why two responding species, represented by MAGs B1A, B1B, and B2, increase with ACA treatment, while the other species of *Muribaculaceae* do not. Annotations of reconstructed genomes suggest that the responders may possess starch degradation capabilities absent in the non-responders.

By including otherwise unannotated genes, genomic comparisons based on OPFs may reflect shared functional potential better than applying previously defined orthologies. Besides the identification of novel gene families, *de novo* homology clustering (37) also enables differentiation of sub-groups not captured by standard annotations. For instance, hypothetical genes annotated as homologs of SusC, SusD, and SusEF, were clustered into 98, 187, and 191 different OPFs respectively. It is plausible that this sub-clustering captures differences in protein structure with importance in oligo- and polysaccharide recognition, import, and binding. Combined with annotation of characterized functional domains, we hypothesize that these clusters more narrowly predict the polysaccharide utilization ranges of uncultured organisms. Testing these predictions will require characterization of the metabolic potential of these genes after obtaining cultivars or through heterologous expression in appropriate hosts.

We examine the three-guild model proposed by Ormerod and colleagues (20) by extending their dimensional reduction approach to the much larger set of *Muribaculaceae* genomes now available. In this analysis, annotations of B1A, B1B, and B2 are not clearly co-located with members of the α-glucan guild, complicating this simple interpretation. Interestingly, a more nuanced analysis involving OPFs and predicted export indicates that B1A and B1B may have polysaccharide utilization potentials distinct from the α-glucan guild previously described. The improved resolution presented by OPF clusters suggests that this more detailed examination might identify specific functions that discriminate responders from non-responders. The approach is bolstered by the phylogenetic and genomic distinction between B2 and both B1A and B1B, reducing the confounding effects of shared evolutionary history.

A detailed analysis of PULs identified multiple loci shared in both B1A and B1B that appear to be adapted to the degradation of starch or related carbohydrates, due to the presence of an OM localized GH13 containing protein (49). Counterintuitively, B2 had no such PUL, suggesting that its response to ACA may result from other enzymatic capabilities. Of particular interest is a PUL encoding proteins with GH97, CBM20, and CBM69 domains, all of which have documented activity on starch (50, 51). While the only OM-localized hydrolase in this PUL is a GH66, and members of this family have characterized activity on the α-1,6 linkages between glucose monomers in dextran (52), it is plausible that this PUL can be repurposed and confers some ability to grow on starch.

Most compellingly, a gene family encoding a SusA homolog was identified in B1A, B1B, and B2 but in none of the non-responders, presenting the best case for a single enzyme that can confer a growth advantage in ACA-treated mice. While it is unclear how expression of this important component of starch utilization might be regulated, given that it is not located in a PUL in any of the responding populations, SusA is important for growth on amylopectin in *B. thetaiotaomicron* (45). Since inhibition by ACA is variable across enzymes (53), it is possible that ACA treatment results in elevated production of dextrin and maltooligosaccharides in the lower guts of mice due to residual α-amylase activity, even at levels sufficient to prohibit host digestion. Periplasmic hydrolysis of these starch breakdown products may be sufficient for increased abundance of these species in ACA-treated mice.

It is notable that of the closely related variants associated with OTU-1, B1B is found at UT and not UM. We previously observed site-specificity of the ACA response of this species, in which OTU-1 did not have a significantly increased abundance in treated mice at UT, while it was the most dramatic change at UM (14). Differences in the functional potential due to differences in gene content of populations found at each of the sites is one possible explanation for this pattern. Alternatively, differences in the occurrence or gene content of other microbial community members could lead to the differential response of OTU-1 across these sites, for instance by contributing to the partial breakdown of larger starch molecules or through resource competition. Notably, ACA appears to be less inhibitory to members of the Firmicutes than the Bacteroidetes (54). Intriguingly, while we do not conjecture a mechanistic link, an ACA-by-site interaction effect on longevity has been previously observed in the mouse colonies sampled here, with male mice at UT showing a larger increase in median longevity with ACA treatment than those at UM (15, 17).

Despite evidence that large differences in gene content can be found between even closely related populations (55, 56), studies reconstructing genomes from metagenomes have just started to consider these pangenome dynamics (57–61). Improvements to software (e.g. binning) and sequencing technologies (e.g. long reads) will increase the accuracy of physiological inferences based on homology and synteny. The discovery of two populations of OTU-1 therefore demonstrates the value of considering pangenome dynamics, and presents a potential explanation for the observed site-specific response of that taxon. The finding that both variants have the same complement of three PULs apparently specializing in starch utilization and the same SusA homolog does not support the hypothesis that differences in starch utilization potential account for these abundance patterns. We did, however, identify numerous differences in the gene content of B1A and B1B, including variant specific loci that may influence the structure and function of the outer surface of the cell. Given the size of these regions, their parallel physiological roles, and their plausible effect on interactions with the host or other microbes (62), we speculate that these variable loci could result in relevant, functional differences between the two populations.

While these results do not establish a mechanistic explanation for differences in the response of B1A and B1B at UM and UT, nor conclusively identify the pathways that enable starch utilization in B2, they do suggest a number of genomic features that likely contribute to previously observed patterns in taxon abundance. Future studies utilizing metatranscriptomic analysis might demonstrate active expression of these genes, or differential expression in mice treated with ACA compared to controls. Likewise, even in the absence of a B2 cultivar, the potential role of its dextran PUL in enrichment under ACA treatment could be tested using available cultivars, like *B. thetaiotaomicron*, that possess a homologous gene cluster.

## Conclusions

In this study we have reconstructed and described genomes representing 8 species in the family *Muribaculaceae* from the mouse fecal microbiome, and have found features that differentiate those bacterial species that respond positively to ACA treatment from those that do not. This analysis suggests that utilization of starch and related polysaccharides enables increased population size in mice treated with ACA—an α-amylase inhibitor. In addition, two distinct genomic variants of one species were identified that differ in functional gene content, potentially explaining site-specific differences in response. By combining observed changes in relative abundance during experimental manipulation with inferred functional gene content, we are able to study mammalian symbionts in the absence of cultured representatives. This sequence-based approach is broadly applicable in microbial ecology and enables improved understanding of *in situ* dynamics within complex microbial communities.

## Methods

### Mouse treatment, sample collection, extraction and sequencing

Mice were bred, housed, and treated as described in (15). Briefly, genetically heterogeneous UM-HET3 mice at each study site were produced by a four-way cross between (BALB/cByJ x C57BL/6J) F1 mothers and (C3H/HeJ x DBA.2J) F1 fathers, as detailed in (63). Mice were fed LabDiet 5LG6 (TestDiet Inc.) from weaning onwards. Starting at 8 months of age, mice randomly assigned to treatment were fed chow with 1,000 ppm ACA (Spectrum Chemical Manufacturing Corporation). Mice were housed 3 males or 4 females to a cage. Colonies were assessed for infectious agents every 3 months, and all tests were negative.

Individual fecal pellets were collected from a single mouse per cage. 16S rRNA gene libraries and metabolite analyses of these samples are as described previously (14). From this collection, a subset of samples was non-randomly selected for metagenomic sequencing in order to test several unrelated hypotheses about SCFA production. Samples were from 54 mice, with at least six treated and control representatives of both males and females at each site.

Fecal samples were slurried with nuclease free water at a 1:10 (w/v) ratio, and most samples were spiked with *Sphingopyxis alaskensis* RB2256 prepared as described previously (14) before DNA extraction and sequencing. Based on alignment to the reference genome, sequenced reads from *S. alaskensis* can be distinguished from all endogenous bacteria in mouse feces. This spike was added as an internal standard to quantify total 16S rRNA gene abundance in a separate study (14). A small number of these were split for both spiked and unspiked samples, which we used to validate this procedure. For each, 150µL of this sample was transferred for extraction using the MoBio PowerMag Microbiome kit. Metagenomic libraries were prepared using standard procedures sequenced on the Illumina HiSeq 400 platform using the v4 paired-end 2×150 bp.

### Assembly, binning, and MAG refinement

Bioinformatic processing of metagenomes was implemented as a Snakemake pipeline (64) run with version 5.18.1. Code and metadata can be obtained at https://doi.org/10.5281/zenodo.4450697.

Raw metagenomic reads were deduplicated using FastUniq (65) version 1.1, adapters trimmed using Scythe (66) version 0.991, and quality trimmed using Sickle (67) version 1.33 to produce processed reads for all downstream analyses. The resulting paired-end reads from all samples were co-assembled into primary contigs using MEGAHIT (68) version 1.2.9. Reads were then mapped back to these contigs with Bowtie2 (69) version 2.4.1, and per-library coverage was estimated for each contig.

For all contigs >1000 bp in length, dimensional reductions built into CONCOCT (70) version 1.1.0 were applied to produce input data for a Gaussian mixture model (GMM) similar to the procedure used by that program for binning. However, unlike CONCOCT—due to computational limitations—the model was trained on just 10% of the input data, sampled randomly, before assigning bins to all contig. While this may have reduced the accuracy of the binning procedure, we believe that subsequent refinement steps mitigated the impact of this decision.

Using 16S rRNA gene libraries described in (14) and processed as in that reference, OTUs were classified taxonomically and relative abundance was calculated. Bins were then recruited to one or more OTUs by calculating a Canonical partial least squares between OTU abundance and bin coverage as implemented in the scikit-learn machine learning library (71) version 0.23.2. For bins recruited to OTUs classified as *Muribaculaceae*, contigs were re-clustered based on coverage across samples. First “trusted contigs” were manually selected which correlated closely with OTU abundance. The mean coverage of these was used to normalize the per-library coverage of all other contigs. Then, using a GMM, groups of contigs were clustered such that the normalized coverage across samples was consistent. These groups were used to inform the manual assignment of contigs to MAGs. Libraries in which MAGs had non-negligible coverage were identified manually and used in subsequent refinements. While clustering contigs associated with OTU-1 a number of groups containing on the order of 10^5^ bp were found with a bimodal coverage distribution, low normalized coverage in a subset of libraries, and normal coverage in others. By this criterion, contigs in these “variable” groups were partitioned into two MAG variants, A and B, with the non-variable contig groups shared by both. To avoid challenges associated with admixture, only libraries that appeared on further inspection to have coverage over just one of the two variants were considered in downstream refinement steps. The mice matching these libraries are highlighted in Fig. 3. None of the other *Muribaculaceae* OTUs described in this study were found during this manual curation process to have genomic variants comparable to B1A and B1B.

For each MAG, the list of contigs and relevant libraries identified in the previous step, as well as the per-library coverage of trusted contigs were used in a final refinement procedure. All read pairs from the relevant libraries with at least one read mapping to the contigs were extracted and used in a single pass of the Pilon assembly refinement tool (72) version 1.23. Regions of these improved contigs were then excised where they had low cosine similarity to the trusted contig coverage, using a 0, 0.5, 0.6, 0.7, 0.8, and 0.9 cosine similarity threshold. Genome completeness and contamination estimates were calculated based on ubiquitous single-copy genes using the program CheckM (73) version 1.1.2. Based on these results, the final refined assembly with the highest completeness and with contamination < 2% was selected; ties were broken using the N50 statistic.

### Reference genomes

In order to compare our 9 novel MAGs to previously reconstructed genomes, we downloaded a total of 98 draft genomes from GenBank, both isolates and MAGs, representing all genomes taxonomically assigned to the *Muribaculaceae* as of September 2020, along with the 30 MAGs described in (20) (BioProject PRJNA313232). For comparison, nucleotide sequences for *Bacteroides thetaiotaomicron* VPI-5482 (GCA_900624795.1), *Bacteroides ovatus* ATCC-8483 (GCA_000154125.1), *Barnesiella viscericola* DSM-18177 (GCA_000512915.1), *Barnesiella intestinihominis* YIT-11860 (GCA_000296465.1), and *Porphyromonas gingivalis* ATCC-33277 (GCA_000010505.1), were also downloaded.

### Genome annotation

All genomes were initially annotated with Prokka (74) version 1.14.6, which uses Prodigal (75) version 2.6.3 for gene finding. Putative protein sequences were additionally annotated with domains from both the dbCAN database (30) release 6 of carbohydrate-active domains and Pfam (76) release 31.0, using HMMER3 (77, 78) version 3.3. In order to maximize the sensitivity of this analysis, these annotations used a bitscore cutoff of 5.0 and no cutoff (any hit), respectively. Protein sequences were also annotated with KO numbers by BLAST using the KEGG database as of March 2018 as the reference and taking the best hit with a maximum E-value of 1e-10.

Lipoproteins were predicted using LipoP (79) version 1.0a and a score cutoff of 5 and a margin cutoff of 2. Lipoproteins with an aspartate at position +2 relative to the cleavage site were labeled as localized to the inner membrane (80, 81). Periplasmic proteins were identified with SignalP (82) version 4.1. Predicted protein sequences from all annotated genomes were first dereplicated using CD-HIT (83, 84) version 4.8.1 at a similarity threshold of 0.99, then locally all-by-all aligned using the DIAMOND (85) version 0.9.31 implementation of the BLASTp algorithm. Each pair was then assigned a similarity value as the bitscore of their best local alignment normalized by the greater of the two self-alignments. This results in a matrix of pairwise scores reflecting the proximity to perfect homology. Scores less than 0.2 were replaced with 0. Clusters were formed using the MCL algorithm (86) version 14-137 with an inflation parameter of 5.

SusCDEF homologs were identified based on relatively relaxed criteria, harnessing OPF assignments, domain predictions, and Prokka annotations to avoid false negatives while maintaining specificity. For each OPF, all KOs assigned to members were collected as plausible KOs for the cluster. Protein sequences in OPF clusters which included K21572 were flagged as putative SusC-homologs, as were sequences directly annotated as such by Prokka. Using a similar approach, proteins in clusters tagged with K21571 or with any of domains PF12771, PF14322, PF12741, PF07980 were identified as putative SusD. Proteins in clusters tagged with K21571, or with either PF14292 or PF16411, were considered SusEF homologs. Putative PULs were identified by “tandem susCD-like pairs”: a SusC-homolog within 5 kbp of a SusD-homolog on the same strand, similar to (87). Of the 121 putative PULs identified across the nine novel MAGs in this study, 103 of them have at least one gene within 10 kbp with homology to a CAZy domain family, suggesting that the majority of our identified PULs may indeed have activity on polysaccharides.

Annotations were compared across all genomes using dimensional reduction (Fig. 1 panels B and C). In order to reduce the effects of redundancy on ordination, highly similar genomes were first clustered by complete-linkage at a 0.8 cosine similarity threshold. Clusters were then replaced by their mean counts before PCA. Finally, the original genomes were projected onto these PCs and visualized.

### Phylogenetics

Predicted amino acid sequences for ORFs from all MAGs and all reference genomes were searched for homology to the TIGRFAM protein clusters (88) release 14.0 using HMMER3. Hits were filtered at the “trusted-cutoff” score threshold defined separately for each protein model. Sequences found in no more than one copy and in at least 95% of genomes were used as taxonomic marker genes for phylogenetic analysis. Marker gene sequences were aligned to their respective models using HMMER3, dropping unaligned residues. Aligned markers were then concatenated for each genome. The concatenated alignment was masked using Gblocks [Castresana2000] version 0.91b using a wrapper script provided at https://github.com/mattb112885/clusterDbAnalysis/blob/master/src/Gblocks_wrapper.py with the default parameters, before estimating an approximate maximum likelihood phylogeny using the FastTree software (89) version 2.1.10 with the default parameters.

## Acknowledgments

We would like to acknowledge the technical support from Randy Strong and others at The University of Texas Health Science Center at San Antonio for initial sample collection. We would also like to thank Nicole Koropatkin for feedback on the manuscript.

## Data Availability

MAGs generated for this study have been deposited to GenBank under the accessions JAEKDF000000000 (B1A), JAEKDG000000000 (B1B), JAEKDH000000000 (B2), JAEKDI000000000 (B3), JAEKDJ000000000 (B4), JAEKDK000000000 (B5), JAEKDL000000000 (B6), JAEKDM000000000 (B7), JAEKDN000000000 (B8). Metagenomic libraries are also available under BioProject PRJNA448009. Code and additional data/metadata needed to reproduce this analysis are available at https://doi.org/10.5281/zenodo.4450697.

## Supplementary Materials

Supplementary Materials are available at https://doi.org/10.5281/zenodo.4450697.

- ***Supplementary Figure S1:*** *Phylogenetic tree identical to Fig. 1 panel A, but with the addition of GenBank accessions for all publicly available genomes*.
- ***Supplementary Figure S2:*** *Phylogenetic tree constructed as in Fig. 1 panel A, but using only the RpoB protein sequence instead of a concatenated gene tree*.
- ***Supplementary Table S1:*** *Annotations, including KO, COG, OPF, Pfam, dbCAN, and localization for all predicted protein coding genes in all genomes analyzed for this study*.
- ***Supplementary Table S2:*** *B1A and B1B gene specificity scores, where the specificity of a gene to the B1A genomic variant is calculated as the ratio of the median normalized coverage in B1B specific libraries versus B1A specific libraries*.
- ***Supplementary Results `build/otu_correlation_and_aca_response.ipynb.html`***: *Assignment of MAGs to OTU labels and responder/non-responder categories*.
- ***Supplementary Results `build/median_relative_mag_coverage.tsv`***: *Normalized MAG coverage data across samples analyzed in this study*.
- ***Supplementary Results `build/otu_relative_abundance.tsv`***: *Relative abundance of Muribaculaceae OTUs across samples analyzed in this study*.
- ***Supplementary Results `build/gh13_families_to_opfs_mapping.ipynb.html`***: *Cross-comparison of sequence membership in GH13 subfamilies and OPFs*.
- ***Supplementary Results `build/gh13_tree.nwk`***: *Estimated phylogeny of GH13 domains from Muribaculaceae, labeled by source genome, OPF, and localization*.
- ***Supplementary Results `build/gtdbtk_classify_summary.tsv`*** *Results from GTDB-tk classify workflow run on the 8 novel MAGs analyzed in this study*.
- ***Supplementary Results `build/B1_inter_strain_comparison.ipynb.html`***: *Analysis and plotting code for coverage and gene content comparisons for B1A and B1B*.
- ***Supplementary Results `build/annotation_ordination.ipynb.html`***: *Analysis and plotting code for annotation and comparative genomics of Muribaculaceae genomes*.
- ***Supplementary Results `build/genome_statistics_table.ipynb.html`***: *Code to compile genome statistics in Tbl. 1*.
- ***Supplementary Results `build/genomic_diagrams.ipynb.html`***: *Code used for automated plotting of PULs for Fig. 2*.
- ***Supplementary Results `build/match_2019_2020_otus.ipynb.html`***: *Analysis code confirming the assignment MAGs in this work to those described in (14)*.
- ***Supplementary Results `build/metabinning_and_strain_variation_muribaculaceae.ipynb.html`***: *Analysis code used to assign contigs, samples, and coverage profiles to the novel MAGs included in this study*.
- ***Supplementary Methods `build/phylogenetic_tree_figure.ipynb.html`***: *Code used to plot annotated species and gene trees*.

