## Supplementary figures and images for "*Muribaculaceae* genomes assembled from metagenomes suggest genetic drivers of differential response to acarbose treatment in mice"

### Supplementary Figure S1

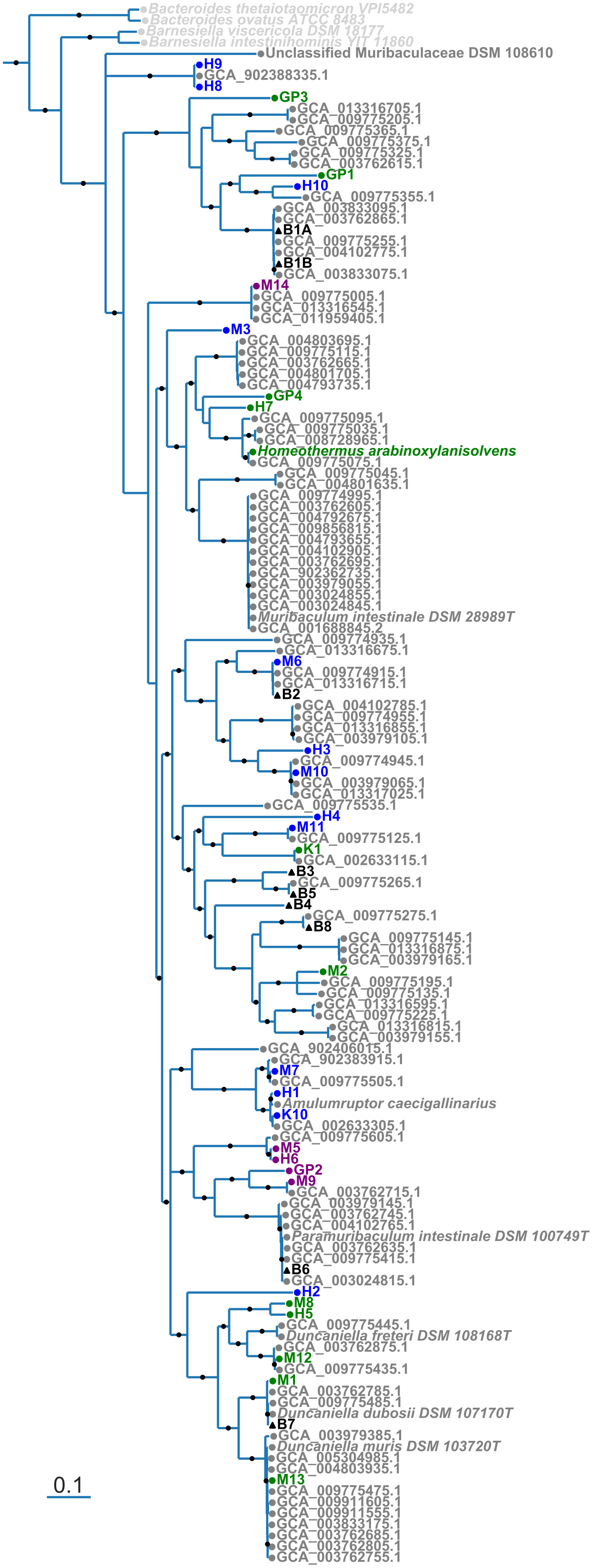

### Supplementary Figure S2

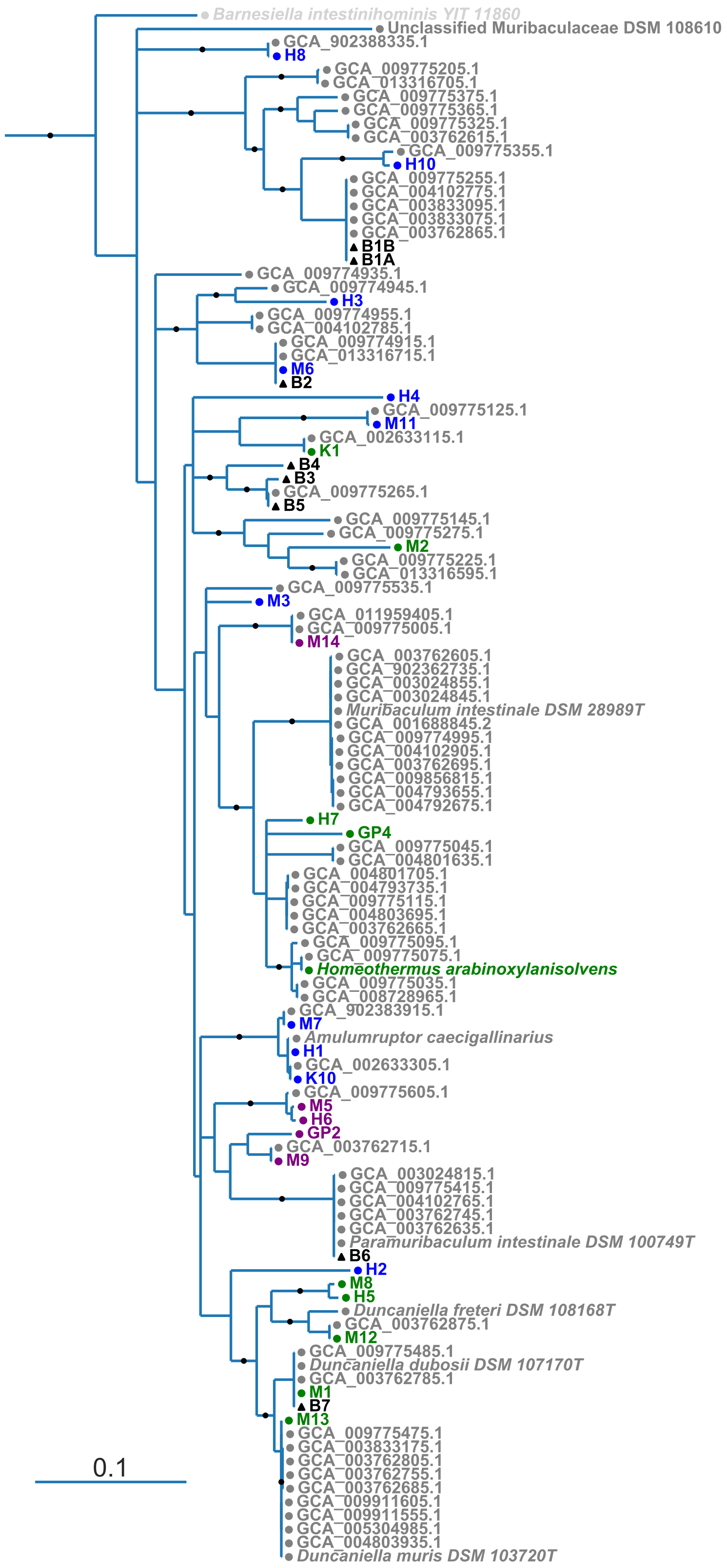
